# Cystinosin deficient rats recapitulate the phenotype of nephropathic cystinosis

**DOI:** 10.1101/2021.06.29.450444

**Authors:** Jennifer A Hollywood, Prasanna K Kallingappa, Pang Yuk Cheung, Renita M Martis, Sree Sreebhavan, Aparajita Chatterjee, Emma J Buckels, Brya G Mathews, Paula M Lewis, Alan J Davidson

## Abstract

**Background:** The lysosomal storage disease cystinosis is caused by mutations in *CTNS*, encoding a cystine transporter, and in its severest form leads to proximal tubule dysfunction followed by kidney failure. Patients receive the drug-based therapy cysteamine from diagnosis. However, despite long-term treatment, cysteamine only slows the progression of end-stage renal disease and a kidney transplant is inevitable. Pre-clinical testing in cystinotic rodents is required to evaluate new therapies; however, the current models are sub-optimal. To solve this problem we generated a new cystinotic rat model.

**Methods:** We utilized CRISPR/Cas9-mediated gene editing to disrupt exon 3 of *Ctns* and measured various parameters over a 12-month time-course including blood and tissue cystine levels, urine and serum electrolytes, and analysed the histopathology and immunohistochemistry of the kidney.

**Results:** *Ctns*^-/-^ rats display hallmarks of cystinosis by 3-6 months of age as seen by a failure to thrive, excessive thirst and urination, cystine accumulation in tissues, corneal cystine crystals, a loss of Lrp2 in proximal tubules and immune cell infiltration. High levels of glucose, calcium, albumin and protein are excreted at 6-months of age, consistent with the onset of Fanconi syndrome, with a progressive diminution of urine urea and creatinine from 9-months of age, indicative of chronic kidney disease. The kidney histology and immunohistochemistry showed proximal tubule atrophy and glomerular damage as well as classic ‘swan neck’ lesions. Overall, *Ctns*^-/-^ rats show a disease progression that more faithfully recapitulates nephropathic cystinosis than existing rodent models.

**Conclusions:** The *Ctns*^-/-^ rat provides an excellent new rodent model of nephropathic cystinosis that is ideally suited for conducting pre-clinical drug testing and a powerful tool to advance cystinosis research.

**Translational Statement:** Good animal models of disease are essential to perform pre-clinical testing of potential new drugs and therapies before they can progress to human clinical trials. Unfortunately, the cystinosis field has been hampered by a lack of suitable animal models that fully recapitulate the disease. We have overcome this by generating a new rat model of cystinosis by CRISPR-Cas9 gene editing. These rodents closely model the human condition in a time-frame that makes them an excellent model ideally suited for pre-clinical drug testing as well as being a powerful tool to advance cystinosis research.

## Introduction

Nephropathic cystinosis is a rare lysosomal storage disease caused by mutations in the *CTNS* gene encoding the cystine transporter, Cystinosin. Defects in this transporter result in the accumulation of cystine within lysosomes of all cells of the body, with insoluble cystine crystals forming in some tissues.^1^ Patients present between 6 to 12-months of age with a loss of essential nutrients in the urine (Fanconi syndrome) and progress to kidney failure before the age of 10 years if untreated.^2^ Widespread cystine accumulation results in a multisystemic disease leading to other symptoms including growth retardation, photophobia due to cystine crystals in the cornea, hypothyroidism, muscle wasting and neurological deterioration. The only available drug for cystinosis is cysteamine, which depletes lysosomal cystine by disulphide bond cleavage, generating a mixed disulphide that exits the lysosome via a lysine transporter.^3^ Cysteamine therapy has many disadvantages, including a strict dosing regimen, sweating odours, and severe gastrointestinal effects. These disadvantages greatly affect drug compliance especially in young adults.^4^ Despite life-long treatment, cysteamine is ineffective against Fanconi Syndrome and only slows the progression of renal damage with patients eventually needing a kidney transplant.^5–7^ Thus, there is a strong clinical need to develop better drug-based therapies for cystinosis.

Animal models are essential for pre-clinical testing of new drug treatments before progressing to clinical trials in humans. Ideally, these models should show a phenotype that closely recapitulates the progression and severity of the human disease. In the case of cystinosis, the first animal model was a mutant mouse generated on a mixed C57BL6/129sv background. Although C57BL6/129sv *Ctns*^-/-^ mice display cystine accumulation and some ocular defects, they lack the proximal tubulopathy and do not undergo renal failure.^8^ A second-generation mouse was backcrossed onto a pure C57BL/6 background and this showed additional features of cystinosis, including Fanconi syndrome, failure to thrive, polydipsia and polyurea.^9^ However, renal failure manifests late in this model, starting at 10-18-months of age, and has a more chronic pathogenesis than that seen in humans.^10–12^ In terms of drug testing, the late onset of disease in the C57BL/6 *Ctns*^-/-^ model may necessitate long term treatment regimens, making the model a challenge to use for pre-clinical studies.

A zebrafish model of cystinosis was developed in 2017 and is characterised by an accumulation of cystine, enlarged lysosomes and proximal tubule dysfunction.^13^ However, classic histopathological features of cystinosis, such as ‘swan neck lesions’ and cystine crystal deposition, are not observed, raising doubts about how well this model recapitulates the human disease. From a drug screening perspective, it is difficult to perform pharmacokinetic analyses on zebrafish and drug delivery in an aquatic environment has a number of challenges.

A genetic linkage analysis that set out to identify the genetic cause of urinary glucose in the Long-Evans Agouti (LEA/Tohm) rat led to the unexpected discovery that this strain has a deletion in the *Ctns* gene. These rats were found to accumulate cystine in their kidneys and display proximal tubular atrophy at ~12 months of age.^14^ However, males of this strain also spontaneously develop non-obese type 2 diabetes, raising the likelihood that additional mutations are present in LEA/Tohm background that could complicate the use of this model for pre-clinical drug testing. Nevertheless, a rat model of cystinosis is highly desirable, as rat physiology more closely resembles that of humans compared to other rodents.^15^

Here, we generated a new rat model of cystinosis using CRISPR/Cas9 to introduce frameshift mutations in the *Ctns* gene in the Sprague Dawley (SD) background. The resulting *Ctns*^-/-^ rats were phenotypically characterised and found to display classic features of cystinosis such as cystine accumulation (including corneal crystals) and kidney pathology (Fanconi syndrome and ‘swan neck’ lesions) that manifested from 3-6 months of age. The close similarity of the *Ctns*^-/-^ rat phenotype to humans with cystinosis, together with the early age of disease onset compared to other rodent models, make *Ctns*^-/-^ rats a valuable new tool to study cystinosis and undertake pre-clinical trials with potential new therapies.

## Materials and Methods

### Animal husbandry and Targeted disruption of *Ctns*

All experimental procedures were approved by the Animal Ethics Committee of the University of Auckland (UoA) in accordance with the New Zealand Animal Welfare Act. The SD [Crl:CD(SD)] IGS rats were supplied by Charles River, USA, and maintained at the Vernon Jensen Unit of the UoA. All animals were kept in temperature (22 ± 2°C), humidity (55 ± 10%) and 12 hr light cycle controlled rooms, with unlimited access to standard pellet food and tap water. The *Ctns* knockout rats were generated using the procedure similar to that described previously (Li et al 2013; Quadros et al. 2017)^16,17^ with some modifications. Recombinant Cas9 protein - Alt-R™ S.p. Cas9 Nuclease 3NLS (Integrated DNA Technologies Pte. Ltd., Singapore) was used and full-length guide (g) RNA targeting exon 3 of *Ctns* (gRNA-ex3 5’-ATCTTTCCAGAATCAACCGTCGG-3’) was produced using the Precision gRNA Synthesis Kit (Thermo Scientific). Online tools RGEN (http://www.rgenome.net/cas-designer/) and COSMID (http://crispr.bme.gatech.edu/)^18,19^ were used to design the target sequence and identify potential off-target sites. Briefly, using forward and reverse overlapping oligonucleotides, the gRNA template was assembled with a T7 promoter in a short ‘one-pot’ PCR reaction. The assembled product was used as the template in an *in vitro* transcription reaction followed by a rapid purification step, yielding gRNA eluted in nuclease-free water. gRNA was quality checked by agarose gel electrophoresis and Nanodrop. The gRNA + Cas9 ribonucleoprotein (ctRNP) complex for microinjection was prepared by diluting the gRNA and Cas9 protein in microinjection buffer to the final concentration of 50 ng/μl and 100 ng/μl, respectively. SD male rats used as sperm source were >10 weeks old, and SD female rats used to obtain oocytes were 4-5 weeks old. The pseudopregnant female SD rats used as embryo recipients were 8-16 weeks old, and the vasectomised SD males used for mating with these females were >10 weeks old. Female SD rats were super-ovulated by IP injection of pregnant mare serum gonadotropin (30 IU; ProSpec, Israel), followed by an injection of human chorionic gonadotropin (25 IU; Sigma-Aldrich, USA) 48 hrs later. Super-ovulated SD females were then mated with male SD rats (1:1) overnight. The day after mating, pronuclear-stage embryos were collected from oviducts of females in M2 medium (Sigma-Aldrich) and transferred into mR1ECM medium (Cosmo Bio Co. Ltd, Japan) until the microinjection. Zygotes with visible pronucleus were microinjected with ctRNP complex both into the pronucleus and the cytoplasm. The microinjected zygotes were cultured in mR1ECM medium until transfer into pseudopregnant females. The embryo transfer was performed either on the same day as microinjection or the next day at the two-cell stage. The microinjected embryos were screened for healthy embryos and transferred into the oviduct of pseudopregnant SD females that were mated with the vasectomised SD males the day before the transfer. The pups were born 20-21 days post embryo transfer. Ear punches were taken from pups at 3 weeks for genotyping and sequencing. Sequencing was also performed at *in silico* identified off-target sites and no indels were discovered. Pups were born at the expected Mendelian ratios indicating that the heterozygous *Ctns* mutation does not interfere with fertility and that the homozygote mutation is not embryonic lethal.

### Cystine Assay

Dissected tissues were immediately placed in 1 ml of 5mM NEM (5 mmol/L in 0.1 M sodium phosphate buffer, pH 7.2) on ice and stored at −80°C until analysis. Tissues were thawed on ice and homogenized in NEM. Protein was precipitated by adding 50 ml sulfosalicylic acid (15% w/v) and samples were centrifuged at 20,000 g for 10 minutes at 4°C. The supernatant was recovered and diluted 1:10 in 0.1% formic acid (due to the extremely high cystine concentrations in homozygote tissues, *Ctns*^-/-^ samples were diluted 1:100). The protein pellet was diluted in 1.5 ml 0.1 M NaOH, and the protein content of the supernatant was determined using Pierce BCA Protein Assay Kit (Thermo Scientific). Cystine content was measured using HPLC-MS/MS as described previously.^20^ A volume of 5 ml of internal standard (20 mM cystine-D4) was added and the sample was pipetted into a glass vial. Chromatographic separation was achieved on a Thermo Scientific Hypercarb column (2.13150 mm; Thermo Scientific) and was maintained at 30°C. Themobile phase consisted of water with 0.01% formic acid and acetonitrile (ACN) with 0.1% formic acid with fast gradient elution at a flow rate of 0.3 ml/min and run time of 5 minutes. The sample volume injected was 4 ml and the autosampler was set at 5°C. Instrument parameters of the mass spectrometer were: gas flow, 6 L/min; gas temperature, 300°C; vaporizer temperature, 250°C; nebulizer, 40 psi; and capillary voltage, 2500 V. Data were acquired and analysed with Agilent Mass Hunter Software. A standard curve was plotted with the observed peak area ratio of analyte to the internal standard against the concentration of the analyte to extract the slope and intercept.

### Plasma and Urine Analysis

Blood was collected in-life from the saphenous vein and placed in heparin containing tubes. Samples were centrifuged at 10,000 g for 10 minutes at 4 °C to obtain plasma. Urine was collected on ice by housing the animals in a metabolic cage for 24 hrs. Water intake and urine volume were recorded. Initial urine analysis was conducted using a Combur^10^ Test as per the manufacturer’s instructions (Roche). In-depth analysis of protein, albumin, glucose, calcium, phosphate, urea and creatinine levels from plasma and urine were assayed on a c331 chemical analyser using commercially available kits, (TP2, ALBT2, GLUC3, CA2, PHOS2, UREAL and CREJ2; Roche). Blood Urea Nitrogen was assayed using Urea nitrogen colorimetric detection kit as per manufacturer’s instructions (Thermo Scientific). The fractional excretion of phosphate (Fe ratio) was calculated using the following formula: FePi (%) = (24-h urine phosphate × serum creatinine)*100/(serum phosphate × 24-h urine creatinine).

### Histological Analysis

Tissues were fixed in 4% paraformaldehyde (PFA) at 4°C for up to 7 days. Tissues were transferred into 70% ethanol and incubated at 4°C. Over the next 2 days, tissues were transferred through a series of 95% and 2 x 100% ethanol, 50:50 ethanol/xylol, 100% xylol, 1 hour each, rocking at room temperature (RT), followed by 50:50 xylol/paraffin at 65°C overnight, and changes of paraffin every 4 hours. After embedding, the blocks were sectioned at 6 μm on a Leica microtome. Sections were air-dried and then stored at 4°C. Sections were stained with haematoxylin and eosin (H&E) and imaged using a Zeiss Axio Imager Z2 microscope. Ten random areas of interest were determined using Image J analysis software macros and histopathological scoring was performed using the following grading criteria: none (0), mild (1-25%), moderate (26-50%) and severe (>50%).^21^ The total number of tubules in each field of view was recorded and assessed for the appearance of proximal tubule atrophy, dilation, basement membrane thickening and epithelial layer effacement. Infiltration of inflammatory cells and glomerular damage were also recorded.

For immunohistochemistry, paraffin sections were deparaffinized at 65°C for 30 minutes, then incubated in two changes of xylene (10 minutes each). Sections were incubated in 3% hydrogen peroxide following antigen retrieval (10 mM sodium citrate buffer plus 0.05% Tween 20, pH 6.0, at 95°C for 30 minutes). After three washes, sections were blocked at RT for at least an hour in blocking solution (Tris-buffered saline (TBS) containing 5% bovine serum albumin (BSA) and 10% normal horse serum with 0.1% Triton X-100). Sections were incubated with primary antibodies; SQSTM1/p62 (1:100) (109012 Abcam); Lamp1 (1:400) (24170 Abcam); Lrp2 (1:500) (110-96417, Novus biologicals); Havcr1 (1:100) (1-76701, Novus biologicals); CD68 (1:100) (MCA-341R, Bio-Rad); CD206 (1:100) (AF2534, R&D systems) in the blocking solution overnight at 4°C in a humidified chamber. After 24 hours, sections were washed three times with 1x TBST (TBS containing 0.1% Triton X-100) and incubated with secondary antibodies; anti-mouse Alexa Fluor 488 (96871 Abcam); Anti-Rabbit Alexa Fluor 594 (96901 Abcam); anti goat Alexa Fluor 594 (SA5-10088 Life technologies) at 1:600 dilution in the blocking solution for 2 hours at RT. Sections were incubated with 10 mg/ml Hoechst 33258 for 5 minutes, washed twice with TBST, and mounted with ProLong Gold (Thermo Scientific) before imaging using a Zeiss LSM710 confocal microscope.

### Image analysis of macrophages

CD68 and CD206 confocal raw images at 20x magnification (approximately 10 random fields per condition) were analysed using ImageJ software. The area of staining of nuclei, green and red positive cells (CD68 and CD206 respectively) were individually quantified using the ImageJ analysis tool and the percentage of green or red staining per cell was calculated for each condition.

### Transmission Electron Microscopy

Tissues were fixed in 2.5% glutaraldehyde and 0.1 M phosphate buffer, pH 7.4, at 4°C and kept in the fixative until processing. Samples were washed three times with 0.1 M phosphate buffer for 10 minutes, then fixed in 1% osmium tetroxide in 0.1 M phosphate buffer for an hour at RT and washed twice in 0.1 M phosphate buffer for 5 minutes. The samples were then dehydrated in a graded series of ethanol washes for 10 minutes each at RT (50%, 70%, 90%, and twice at 100%), followed by two propylene oxide washes for 10 minutes at RT. The samples were then infiltrated with a graded series of propylene oxide/ resin mix (2:1, 1:1, 1:2) for 30 minutes each, before being embedded in freshly-prepared pure resin overnight. The next day, the samples were placed into moulds and polymerized at 60 °C for 48 hours. All washes were performed on a rocker. Sectioned samples were imaged using a Tecnai G2 Spirit Twin transmission electron microscope.

### Immunoblotting

Tissues were homogenised in 500 μL of ice-cold radioimmunoprecipitation assay buffer (Thermo Scientific) supplemented with protease (cOmplete Mini; Roche) and phosphatase inhibitors (PhosSTOP; Roche). Samples were centrifuged at 12,000 g for 10 minutes at 4°C. Protein content of the supernatant was determined using the Pierce BCA Protein Assay Kit (Thermo Scientific). Equal amounts of protein were boiled in Laemmli buffer at 95°C for 10 minutes. A total of 30 μg of protein was separated by SDS-PAGE and transferred to nitrocellulose membranes (Bio-Rad) using the semidry Trans-Blot Turbo device (Bio-Rad). Membranes were blocked in 2% fish skin gelatine (Sigma-Aldrich) in TBST for 1 hour at RT and probed using specific antibodies for LC3BII (1:1000; 3868 Cell Signalling Technologies), SQSTM1/p62 (1:1000; 109012 Abcam) and ACTB (1:20,000; A2228 Sigma-Aldrich). Primary antibodies were incubated overnight at 4°C with gentle agitation. The next morning, membranes were probed with either anti-rabbit or anti-mouse linked to horseradish peroxidase secondary antibodies (1:20,000 dilution) for 1 hour at RT. Membranes were visualised using enhanced chemiluminescence reagent (ECL Select Kit; GE HealthCare) and chemiluminescent signals were captured using the ChemiDoc image device (Bio-Rad). Densitometry analysis of protein bands was quantified using ImageJ software (National Institutes of Health, Bethesda, MD). The intensity of each band was recorded relative to a pooled control sample run on each gel.

### Behavioural testing

Behavioural tests were carried out on rats that were 9 and 12 months old. Rats were conditioned prior to testing. For the open field test, rats were placed in an enclosure (40 by 40 cm) for a 15 min period. Activity was recorded using a camera suspended above the enclosure and data was analysed using Ethovision XP version 12.Total distance moved, time spent in the inner and outer area and the number of rearing behaviours were recorded.

### Ocular and Bone examination

Corneas of 2, 3 and 9-month-old *Ctns*^-/-^ and wild-type anesthetized rats were examined using the Micron IV slit lamp attachment (Phoenix Research Lab, California, USA). Body temperature was maintained with the use of a heating pad and artificial tears (Systane Ultra; Alcon, Geneva, Switzerland) were instilled regularly to keep the cornea hydrated at all times. X-ray examination were taken of the entire skeleton on 12 and 18-month old anesthetized rats with an Ami-HTX instrument (Spectral Instruments Imaging, Tucson, Arizona) using the high resolution and high power (40kV, 200uA) settings.

### Microcomputed tomography (microCT)

Soft tissue was removed and tibiae were fixed in 70% ethanol at 4°C. The distal end of tibiae were scanned using a Skyscan 1172 microCT scanner (Bruker, Belgium) as previously described.^22^ The X-ray voltage and amperage were 80 kV and 124 μA, and a 1 mm aluminium filter was used. Images were acquired with an isotropic voxel size of 12 μm, with 180° of rotation, and a rotation step of 0.44°. After standardised reconstruction using NRecon software (Bruker, version 1.6.9.18), the datasets were analysed using CTAn software (Bruker, version 1.18.8.0). Standardised parameters of trabecular and cortical bone microstructure were measured.^23^ The trabecular region of interest was 1.07 mm and 0.95 mm distal to the growth plate in females and males respectively, and extended 3.58 mm in the distal direction. The cortical region of interest was 9.54 mm distal to the growth plate, and extended 2.38 mm in the distal direction in both sexes.

### Statistical analysis

Data are presented as the mean ± SEM. GraphPad PRISM software version 7 (GraphPad Software) was used for all statistical analyses. The statistical significance of the differences between two groups was calculated using an unpaired t-test. For two variables, two-way ANOVA was used. A P value <0.05 was considered to be statistically significant.

## Results

### Generation of knockout rats

Using CRISPR/Cas9-mediated gene editing of embryos from SD rats, we generated three independent *Ctns* knockout (KO) lines (allelic designations *Ctns^E3*N-2Vjupk/Vju^, Ctns^E3*N-3Vjupk/Vju^, Ctns^E3*N-4Vjupk/Vju^*) with disruptions in exon 3 of the *Ctns* gene (Figure 1A). These founders possessed 2 (*Ctns^E3*N-2Vjupk/Vju^*) or 8 (*Ctns^E3*N-3Vjupk/Vju^*) bp insertions or 7 (*Ctns^E3*N-4Vjupk/Vju^*) bp deletions, respectively, which result in frameshifts and premature stop codons that cause early termination of translation and are predicted to be complete loss-of-function alleles (Figure 1B-C). The three founder females were back-crossed with wild-type SD males to establish experimental litters following two rounds of breeding. Heterozygous animals were inbred to generate KO homozygotes and wild type littermates and phenotypically characterised (described in detail below). As no phenotypic differences were observed between the three founder lines, the data from these animals (herein collectively referred to as *Ctns*^-/-^ animals) were pooled for analysis.

**Figure 1:**
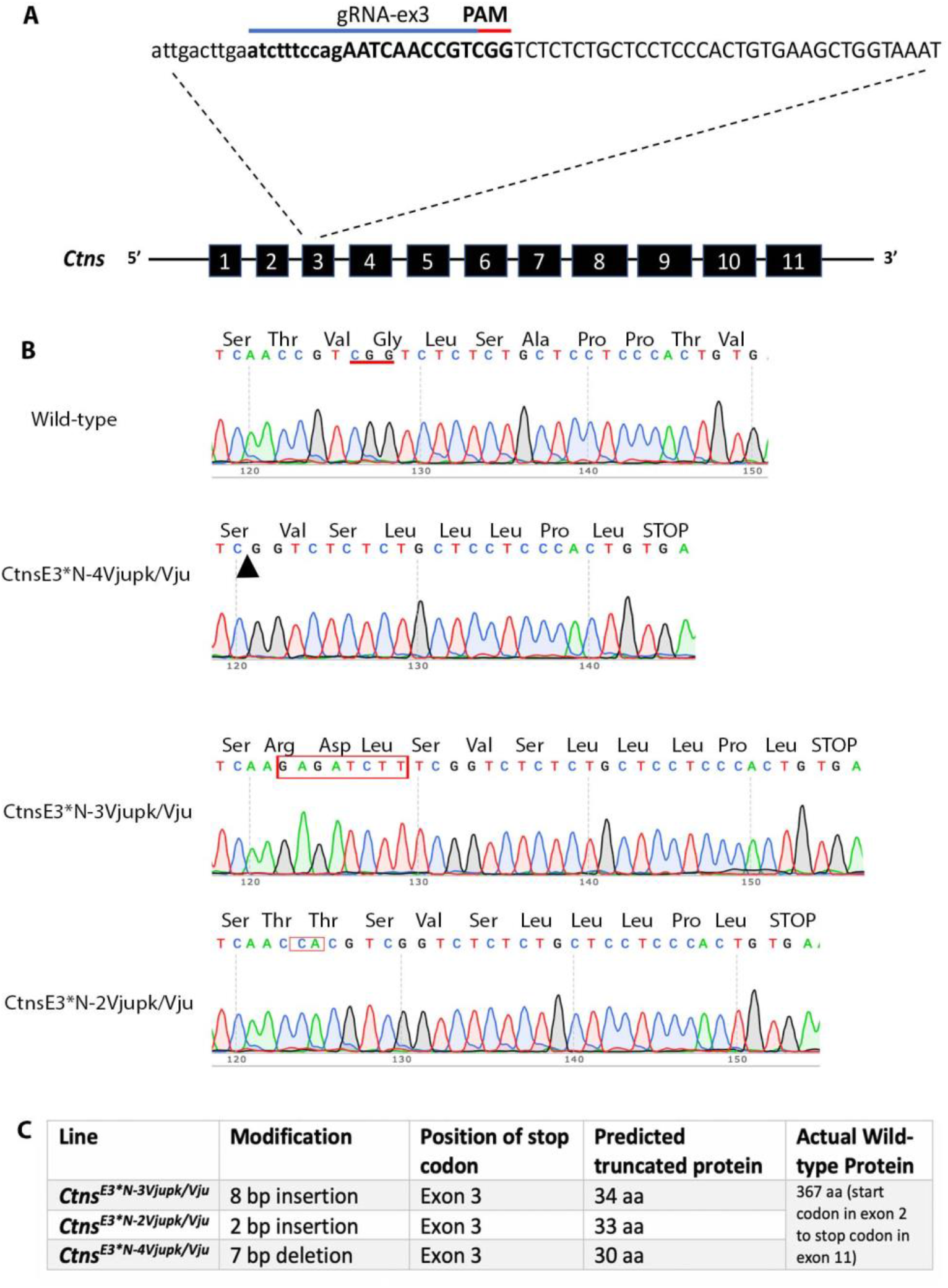
Generation of Cystinosin knock-out (*Ctns*^-/-^) rats. **A)** Schematic overview of the CRISPR-based strategy to disrupt exon 3 of the *Ctns* gene in Sprague Dawley rat embryos. **B)** Sanger sequencing chromatogram with accompanying amino acids shows resulting sequence in *Ctns* Knockout (KO) founders. Position of deletion indicated by black arrow and insertions are boxed in red. **C)** Table indicating modifications and predicted protein. PAM, protospacer adjacent motif.

### *Ctns*^-/-^ rats exhibit failure to thrive

Weekly weighing of *Ctns*^-/-^ rats beginning at weaning (3 weeks) up until 52 weeks of age showed that both male and female *Ctns*^-/-^ rats weigh less than littermate wild-type (WT) controls starting from 16 weeks of age. This weight difference became progressively worse over time such that by 52 weeks of age, *Ctns*^-/-^ male rats weighed on average 25% less than controls while females weighed on average 37% less than controls (Figure 2A and B).

**Figure 2:**
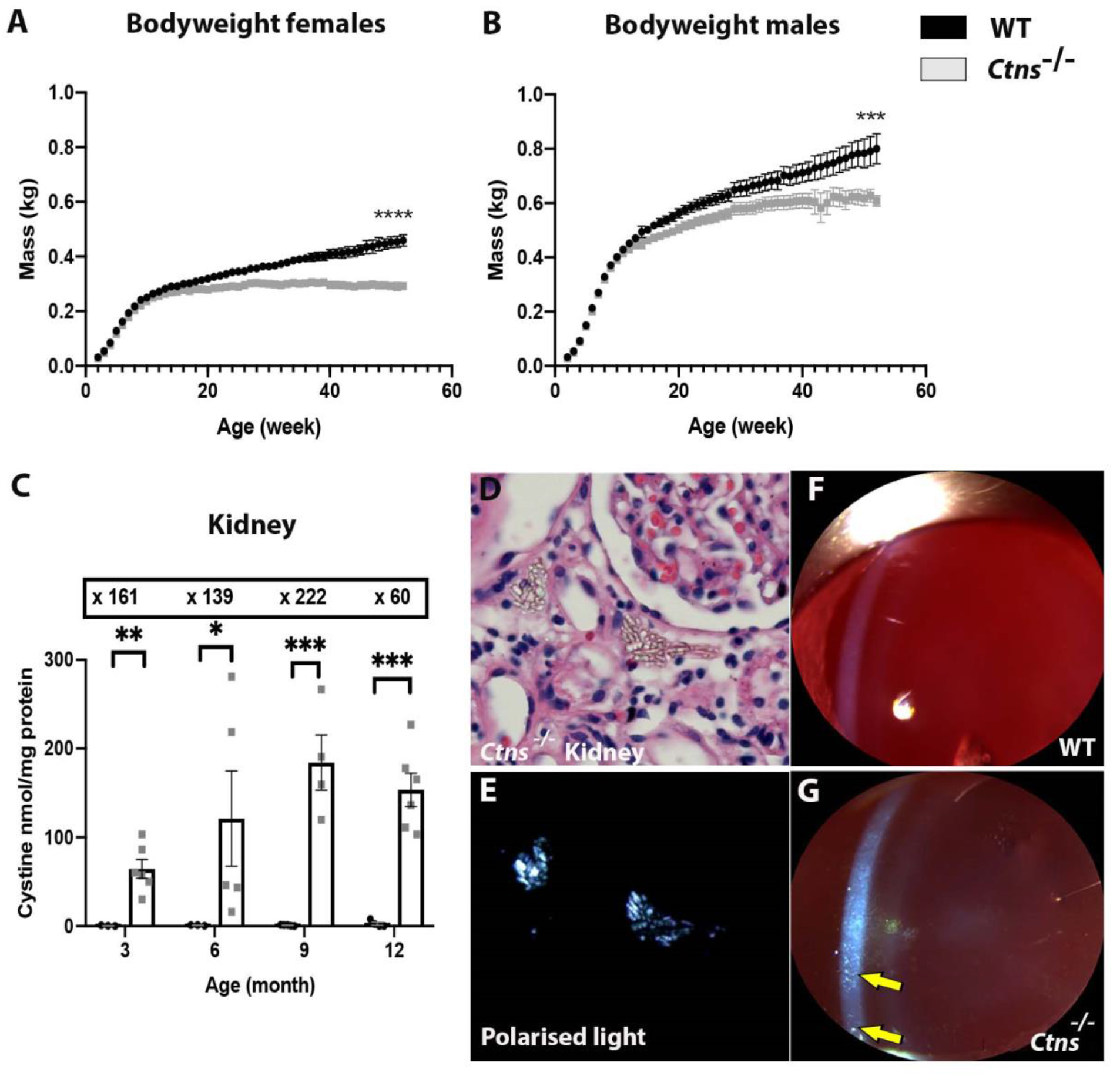
*Ctns*^-/-^ rats exhibit failure to thrive and accumulate cystine. **A, B)** Weight gain (kg) in wild-type (WT) (black) and *Ctns*^-/-^ (grey) female and male rats over the course of 52 weeks. Two-way ANOVA performed. ***P<0.001, ****P<0.0001. n = 38 WT and 41 *Ctns*^-/-^. **C)** Amount of cystine (nmol/mg of protein) in WT and *Ctns*^-/-^ rat kidney at different timepoints. Boxed values indicate fold-change compared to WT at same timepoint. Two-way ANOVA performed. *P<0.05, **P<0.01, ***P<0.001. n = 5 WT and 6 *Ctns*^-/-^ per group. **D)** H & E-stained kidney section from 6-month-old *Ctns*^-/-^ rat showing cystine crystals. **E)** Same section visualised using polarised light. **F)** Slit lamp examination of 3-month-old WT eye showing clear cornea. **G)** Slit lamp examination of 3-month-old *Ctns*^-/-^ displaying cystine crystals in the cornea, yellow arrows. n = 4 WT and 10 *Ctns*^-/-^.

### *Ctns*^-/-^ rats accumulate cystine and deposit crystals

We next measured the intracellular cystine content of the kidney, spleen, heart, soleus muscle, liver, lung and brain from *Ctns*^-/-^ and WT littermate control rats at 3, 6, 9, and 12-months by HPLC-MS/MS. We found that *Ctns*^-/-^ rats show elevated levels of cystine in all tissues examined compared to littermate controls, with the highest observed in the kidney (183 ± 31.10 vs 0.83 ± 0.26 nmol/mg of protein, respectively) at 9-months of age (Figure 2C, Supplementary Figure 1A-F and Table 1). To directly visualise cystine crystals, sections of tissues were examined by polarised light microscopy. This revealed the deposition of numerous crystals in the kidney, muscle, liver and heart starting from 3-months of age (Figure 2D and 2E, and data not shown). The presence of corneal crystals was assessed by slit lamp analysis in 2, 3 and 9-month old *Ctns*^-/-^ and control rats with a slit lamp biomicroscope. While no crystals were detected at the 2-month timepoint, they were readily visible in the corneas of 3 and 9-month old *Ctns*^-/-^ rats. No crystals were noted in the corneas of control animals (Figure 2F and 2G).

**Table 1:** Cystine levels in tissues of wild-type and Ctns^-/-^ rats (nmol/mg of protein)

| Tissue | Genotype | 3 month | 6 month | 9 month | 12 month |
| --- | --- | --- | --- | --- | --- |
| Kidney | WT | 0.409 ± 0.23 | 0.87 ± 0.29 | 0.83 ± 0.26 | 2.56 ± 1.8 |
| | $Ctns^{-/-}$ | 64.5 ± 10.7 | 121.1 ± 53.8 | 184 ± 31.10 | 153.5 ± 18.92 |
| Liver | WT | 0.27 ± 0.16 | 0.31 ± 0.06 | 0.35 ± 0.14 | 0.36 ± 0.10 |
| | $Ctns^{-/-}$ | 4.5 ± 1.01 | 16.8 ± 4.97 | 30.9 ± 6.63 | 18.32 ± 3.43 |
| Lung | WT | 0.55 ± 0.33 | 0.74 ± 0.30 | 0.46 ± 0.19 | 0.33 ± 0.1 |
| | $Ctns^{-/-}$ | 12.4 ± 3.21 | 24.6 ± 10.51 | 46.6 ± 11.25 | 61.3 ± 11.82 |
| Spleen | WT | 0.99 ± 0.36 | 3.67 ± 1.75 | 3.22 ± 2.05 | 0.46 ± 0.20 |
| | $Ctns^{-/-}$ | 161 ± 17.58 | 274 ± 30.9 | 297 ± 60.27 | 373.28 ± 94.6 |
| Brain | WT | 0.14 ± 0.04 | 0.26 ± 0.06 | 0.18 ± 0.04 | 0.33 ± 0.12 |
| | $Ctns^{-/-}$ | 2.1 ± 0.18 | 1.95 ± 0.18 | 5.16 ± 1.65 | 4.42 ± 0.27 |
| Muscle | WT | 0.22 ± 0.01 | 0.82 ± 0.23 | 0.45 ± 0.12 | 0.33 ± 0.01 |
| | $Ctns^{-/-}$ | 6.88 ± 1.19 | 4.04 ± 0.86 | 5.25 ± 0.94 | 9.51 ± 1.15 |
| Heart | WT | 0.19 ± 0.11 | 0.27 ± 0.03 | 0.82 ± 0.39 | 0.78 ± 0.36 |
| | $Ctns^{-/-}$ | 17.6 ± 4.4 | 27.2 ± 8.6 | 230 ± 60.28 | 286.5 ± 83.69 |

### Ctns^-/-^ rats aged 3-6 months show Fanconi syndrome and renal damage

To determine if *Ctns*^-/-^ rats develop a Fanconi-like syndrome we collected urine samples for 24-hours each month and analysed for disturbed water, glucose, calcium, proteins and phosphate excretion. Signs of polyurea and polydipsia in *Ctns*^-/-^ rats were observed from 3-months of age with severity worsening with age (Figure 3A and B). An analysis of urine composition (normalised to urine output) showed a statistically significant increase in daily excretion of total protein, glucose, calcium and albumin at 6-months of age, while an increase in phosphate was detected at 9-months of age (Figure 3C-G and Table 2). Urea and creatinine excretion were significantly decreased at 9-months of age, indicative of compromised kidney function (Figure 3H and 3I). Combur^10^ urinary reagent strip analysis of monthly urine excretion detected significantly higher levels of leukocytes in the urine of *Ctns*^-/-^ rats from 2-months of age, indicative of an inflammatory reaction in the kidney (Figure 3J). Consistent with decreased urine creatinine excretion, plasma creatinine levels were significantly increased in *Ctns*^-/-^ animals at 9-months of age compared to wild-type littermate controls (Figure 3K). Despite these signs of kidney dysfunction, a significant difference in plasma urea and blood urea nitrogen was not detected in *Ctns*^-/-^ rats until 17-months of age (Figure 3L and 3O). No difference in plasma phosphate was found at any timepoint compared to control animals (Figure 3M). To assess nephron stress, we measured the fractional excretion of phosphate (the amount of phosphate filtered by the glomerulus that is excreted into urine) revealing a significant increase in *Ctns*^-/-^ rats at 9-months of age with a further increase at 12-months of age, compared to control animals (Figure 3N and Table 2).^24,25^

**Figure 3.**
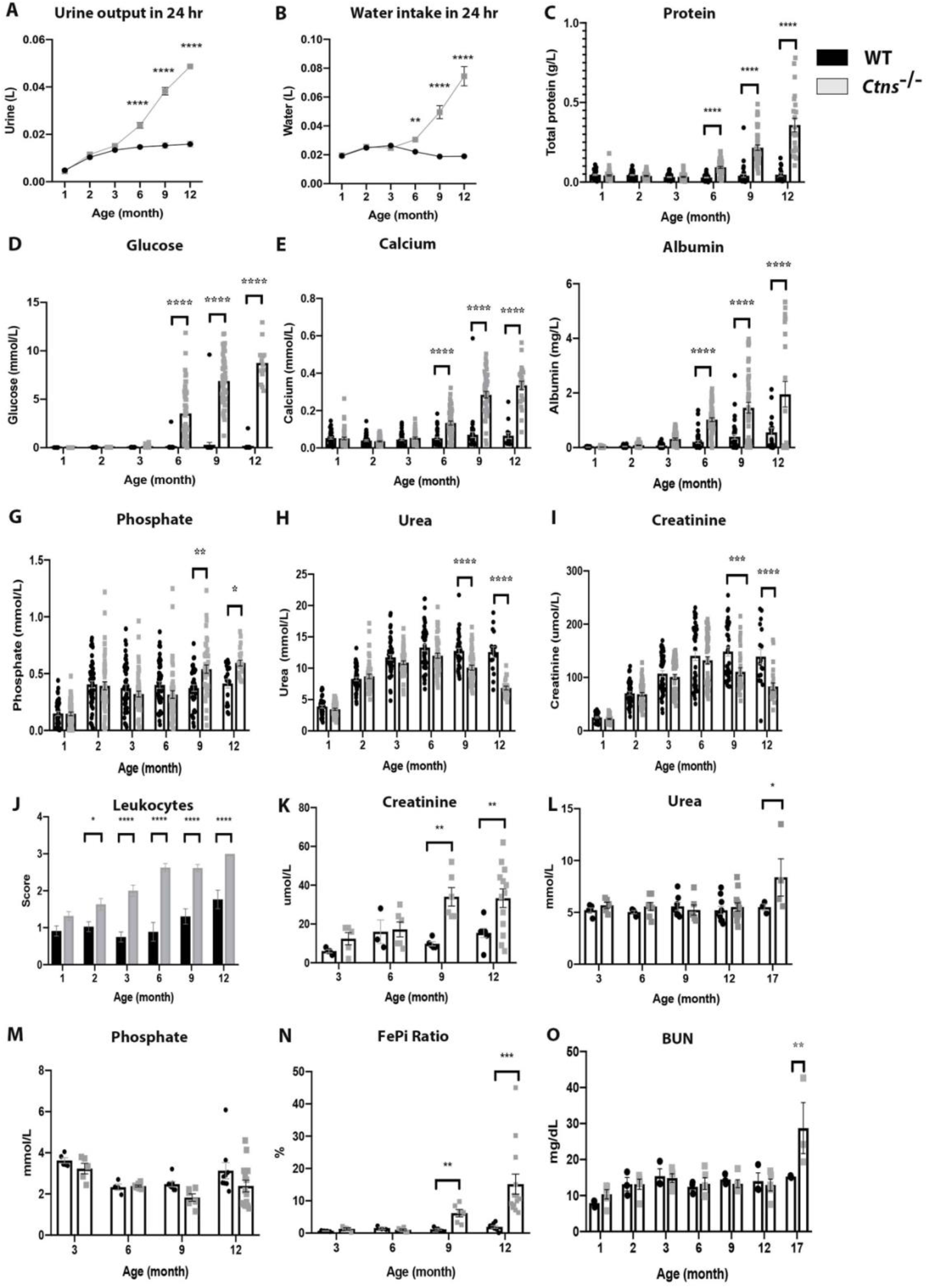
*Ctns*^-/-^ rats aged 3-6 months show clinical Fanconi syndrome. **A)** Volume of urine excreted in 24 hrs in WT and *Ctns*^-/-^ rats. **B)** Volume of water ingested over 24 hrs in WT and *Ctns*^-/-^ rats. Amount of; **C)** protein (g/L), **D)** glucose (mmol/L), **E)** calcium (mmol/L), **F)** albumin (mg/L), **G)** phosphate (mmol/L), **H)** urea (mmol/L) and **I)** creatinine (μmol/L) present in the urine of WT and *Ctns*^-/-^ rats. Two-way ANOVA performed. *P<0.05, **P<0.01, ***P<0.001, ****P<0.0001. n =45 WT and 52 *Ctns*^-/-^. **J)** Amount of Leukocytes in the urine; a score of 0 = normal, 1 = 10-25 leu/μl, 2 = 75 leu/μl, 3 = 500 leu/μl. Two-way ANOVA performed. *P<0.05, ****P<0.0001. n = 45 WT and 40 *Ctns*^-/-^. Amount of **K)** creatinine (μmol/L), **L)** urea (mmol/L) and **M)** phosphate (mmol/L) present in the plasma of WT and *Ctns*^-/-^ rats. Two-way ANOVA performed. *P<0.05, **P<0.01. n= 8 WT and 15 *Ctns*^-/-^. **N)** Graph displaying the ratio of fractional excretion of phosphate (%). Two-way ANOVA performed. **P<0.01, ***P<0.001. n= 26 WT and 30 *Ctns*^-/-^. **O)** Amount of blood urea nitrogen (mg/dL). Two-way ANOVA performed. **P<0.01. n = 3 WT and 5 *Ctns*^-/-^.

**Table 2:** Urine and plasma values in wild-type and Ctns^-/-^ rats

| Urine values | Genotype | 1 month | 2 month | 3 month | 6 month | 9 month | 12 month |
| --- | --- | --- | --- | --- | --- | --- | --- |
| Protein (g/L) | WT | 0.046 ± 0.004 | 0.044 ± 0.003 | 0.032 ± 0.003 | 0.029 ± 0.003 | 0.04 ± 0.01 | 0.047 ± 0.01 |
| | $Ctns^{-/-}$ | 0.042 ± 0.004 | 0.04 ± 0.003 | 0.033 ± 0.003 | 0.093 ± 0.006 | 0.217 ± 0.02 | 0.36 ± 0.042 |
| Glucose (mmol/L) | WT | 0.015 ± 0.001 | 0.034 ± 0.002 | 0.035 ± 0.002 | 0.095 ± 0.06 | 0.291 ± 0.26 | 0.144 ± 0.12 |
| | $Ctns^{-/-}$ | 0.017 ± 0.001 | 0.038 ± 0.002 | 0.097 ± 0.016 | 3.53 ± 0.38 | 6.88 ± 0.33 | 8.74 ± 0.53 |
| Calcium (mmol/L) | WT | 0.053 ± 0.005 | 0.04 ± 0.004 | 0.045 ± 0.005 | 0.051 ± 0.006 | 0.07 ± 0.015 | 0.066 ± 0.015 |
| | $Ctns^{-/-}$ | 0.051 ± 0.007 | 0.035 ± 0.003 | 0.054 ± 0.005 | 0.134 ± 0.011 | 0.285 ± 0.02 | 0.335 ± 0.022 |
| Albumin (mg/L) | WT | 0.014 ± 0.002 | 0.041 ± 0.006 | 0.047 ± 0.009 | 0.206 ± 0.05 | 0.398 ± 0.09 | 0.56 ± 0.16 |
| | $Ctns^{-/-}$ | 0.016 ± 0.002 | 0.083 ± 0.01 | 0.313 ± 0.033 | 1.022 ± 0.07 | 1.46 ± 0.19 | 1.95 ± 0.47 |
| Phosphate (mmol/L) | WT | 0.150 ± 0.02 | 0.405 ± 0.03 | 0.374 ± 0.03 | 0.401 ± 0.03 | 0.37 ± 0.03 | 0.41 ± 0.04 |
| | $Ctns^{-/-}$ | 0.146 ± 0.02 | 0.394 ± 0.03 | 0.032 ± 0.03 | 0.316 ± 0.03 | 0.54 ± 0.03 | 0.59 ± 0.03 |
| Urea (mmol/L) | WT | 3.88 ± 0.23 | 8.3 ± 0.27 | 11.75 ± 0.43 | 13.31 ± 0.54 | 12.75 ± 0.44 | 12.56 ± 0.84 |
| | $Ctns^{-/-}$ | 3.45 ± 0.16 | 8.7 ± 0.35 | 10.89 ± 0.3 | 11.9 ± 0.41 | 10.1 ± 0.4 | 6.84 ± 0.29 |
| Creatinine (μmol/L) | WT | 24.9 ± 1.4 | 70.12 ± 3.3 | 107.14 ± 5.5 | 140.68 ± 8.9 | 148.5 ± 8.4 | 139.2 ± 13.93 |
| | $Ctns^{-/-}$ | 21.99 ± 1.0 | 68.43 ± 3.24 | 100.9 ± 4.74 | 132.12 ± 6.75 | 136.2 ± 8.12 | 82.94 ± 6.7 |

| Serum values | Genotype | 3 month | 6 month | 9 month | 12 month | 17 month |
| --- | --- | --- | --- | --- | --- | --- |
| **Creatinine (μmol/L)** | **WT** | 6 ± 0.82 | 16 ± 6.2 | 9.8 ± 1.12 | 15.5 ± 2.4 | - |
|  | *Ctns*<sup>-/-</sup> | 12.4 ± 3.2 | 17.2 ± 3.8 | 34 ± 4.8 | 33.3 ± 4.8 | - |
| **Urea (mmol/L)** | **WT** | 5.2 ± 0.3 | 5.02 ± 0.15 | 5.6 ± 0.4 | 5.2 ± 0.4 | 5.5 ± 0.3 |
|  | *Ctns*<sup>-/-</sup> | 5.7 ± 0.3 | 5.6 ± 0.40 | 5.2 ± 0.50 | 5.5 ± 0.40 | 8.4 ± 1.8 |
| **Phosphate (mmol/L)** | **WT** | 3.6 ± 0.13 | 2.3 ± 0.15 | 2.48 ± 0.13 | 3.1 ± 0.40 | - |
|  | *Ctns*<sup>-/-</sup> | 3.2 ± 0.26 | 2.4 ± 0.05 | 1.8 ± 0.18 | 2.4 ± 0.28 | - |
| **FePi ratio (%)** | **WT** | 0.64 ± 0.1 | 1.55 ± 0.35 | 1.2 ± 0.24 | 1.9 ± 0.40 | - |
|  | *Ctns*<sup>-/-</sup> | 1.3 ± 0.30 | 1 ± 0.25 | 6.2 ± 1.20 | 15.1 ± 3.10 | - |
|  | **Genotype** | **3 month** | **6 month** | **9 month** | **12 month** | **17 month** |
| **BUN (mg/dL)** | **WT** | 12.3 ± 3.4 | 10.8 ± 1.7 | 13.1 ± 1.5 | 13.5 ± 1.7 | 15.3 ± 0.3 |
|  | *Ctns*<sup>-/-</sup> | 14.9 ± 1.2 | 13.3 ± 1.6 | 13.3 ± 1.1 | 13 ± 1.6 | 28.7 ± 7 |

### *Ctns*^-/-^ rats display glomerular and proximal tubule lesions

Histological analysis of *Ctns*^-/-^ and control kidneys was performed to identify cellular and structural lesions. At 3-months of age, *Ctns*^-/-^ rats showed mild signs of proximal tubule atrophy, basement membrane thickening and tubule dilation and effacement in the region of the superficial cortex (Table 3). At this time, we observed the occasional ‘swan neck’ lesion, the hallmark histological feature seen in human cystinotic kidneys (5% of glomeruli observed, n=6/117; Figure 4B and 4F). We also noted mild renal corpuscle abnormalities that were characterised by glomerular hyper-segmentation and/or shrinkage (Table 3). At 6-months of age, the glomerular and proximal tubule lesions were more moderate with prominent thickening of the basement membrane in places (orange arrow in Figure 4C) as well as areas of inflammatory infiltration (Figure 4G). Swan neck lesions were seen in 9% of glomeruli (n=4/46; yellow arrow in Figure 4C). The above histopathology was even more severe at 9-months of age, with considerable tubular atrophy and dilation (yellow arrows in Figure 4D), and casts in the lumens of tubules at the corticomedullary junction (Figure 4H). At 12-months of age, multiple large foci of severely atrophic tubules were observed. Most of the proximal tubules were unrecognisable with some filled with protein casts (blue arrow in Figure 4E). Glomerular tufts were largely shrunken and/or sclerotic (n=67/89; yellow arrow in Figure 4E and 4I). We observed the presence of multinucleated podocytes and tubule cells at 9- and 12-months of age (Figure 4K and 4L), which is also frequently seen in cystinotic patients. By contrast, we did not detect any of these abnormalities in kidney sections from control animals at any timepoint (Figure 4A and 4J). The markedly different histologies of cystinotic and control kidneys was also reflected at a gross level, with the dissected kidneys from *Ctns*^-/-^ rats being much paler and larger compared to WT kidneys from 6-months of age onwards and being most prominent at 12-months of age (Figure 4M).

**Figure 4.**
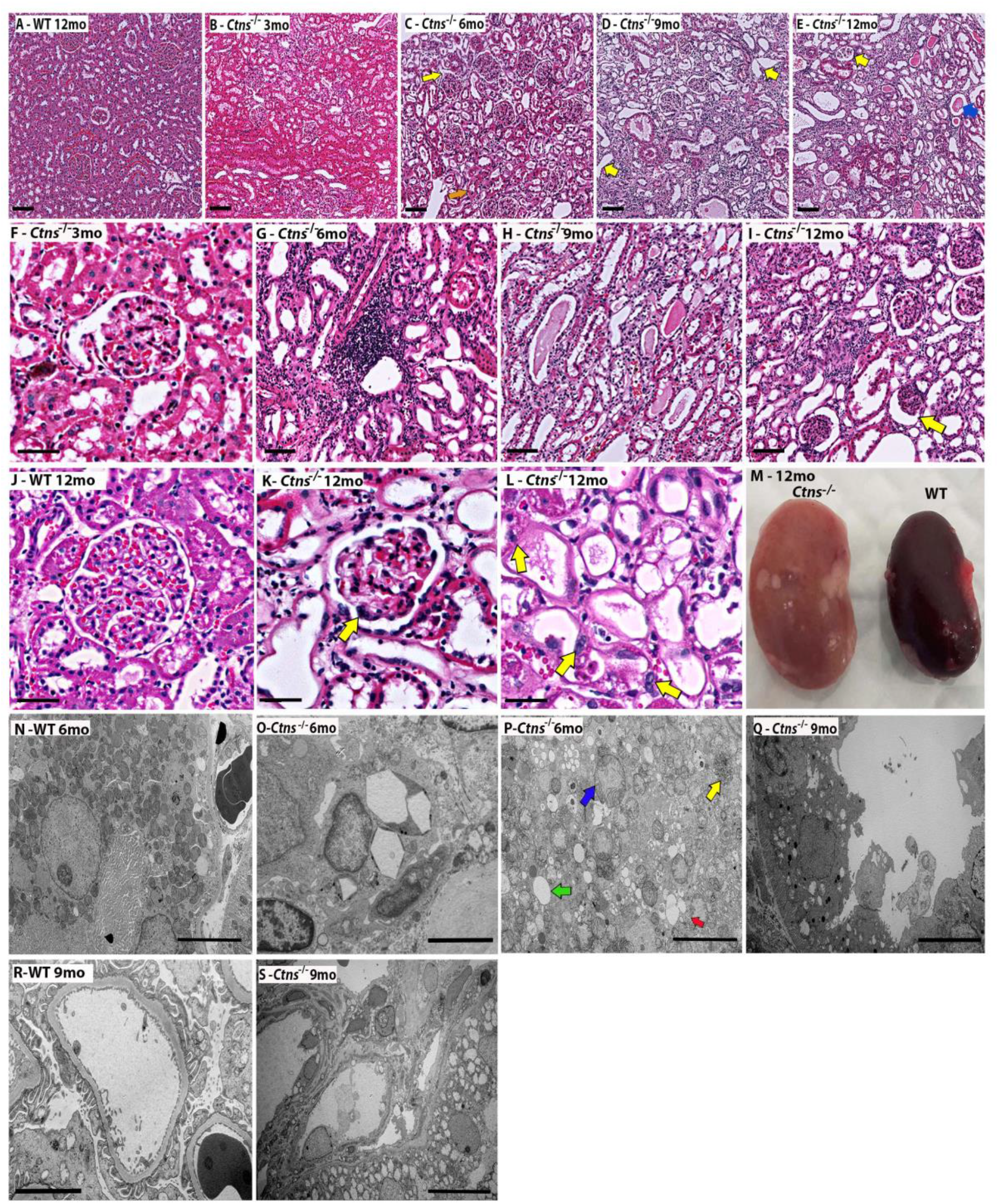
*Ctns*^-/-^ rats display proximal tubule atrophy and histological lesions. **A-L)** Representative H&E stained kidney sections of A) WT and B-E) *Ctns*^-/-^ rats at timepoints indicated. **F)** Swan neck lesion in 3-month-old *Ctns*^-/-^ rat. **G)** Area of inflammation in 6-month-old *Ctns*^-/-^ rat. **H)** Tubular casts 9-month-old *Ctns*^-/-^ rat. **I)** Abnormal shrunken glomerulus, yellow arrow, in 12-month-old *Ctns*^-/-^ rat. **J)** Healthy glomerulus in WT rat. Multinucleated **K)** podocytes and **L)** tubules, yellow arrows, in 12-month-old *Ctns*^-/-^ rat. Scale bars 100 μm (A-E), 50 μm (F-L). **M)** 12-month-old *Ctns*^-/-^ and WT whole kidney at dissection. Transmission electron micrograph of **N)** WT proximal tubule, **O)** cystine crystals in the lysosomes of 6-month-old *Ctns*^-/-^ rat, **P)** enlarged mitochondria and vesicles, **Q)** degenerating proximal tubule, **R)** healthy podocyte in WT and **S)** damaged podocyte in 9-month-old *Ctns*^-/-^ rat. Scale bars, 5 μm in (N, P, Q and S) and 2 μm in (O and R).

**Table 3:** Histological Grading: none (0), mild (1-25%), moderate (26-50%) and severe (>50%)

|  | 3 month | 6 month | 9 month | 12 month |
| --- | --- | --- | --- | --- |
| **Proximal Tubule atrophy** | 20% mild | 45% moderate | 61% severe | 56% severe |
| **BM thickening** | 21% mild | 33% moderate | 38% moderate | 47% moderate |
| **Tubule dilation** | 10% mild | 18% mild | 54% severe | 42% moderate |
| **Epithelial effacement** | 8.5% mild | 30% moderate | 31% moderate | 34% moderate |
| **Hyper-segmented glomeruli** | 29/117 | 6/46 | 26/69 | 46/89 |
| **Glomerulus shrinkage** | 12/117 | 4/46 | 16/69 | 21/89 |

Ultrastructural analysis of kidney sections from *Ctns*^-/-^ rats at 6-months of age showed the presence of intracellular cystine crystals with an angular appearance that were usually located within lysosomes (Figure 4O). We also noted the increased presence of swollen mitochondria (blue arrow), enlarged empty vesicles/lysosomes (green arrow), multivesicular bodies (yellow arrow) and double membraned autophagosomes (red arrow) in *Ctns*^-/-^ cells compared to the healthy control cells (Figure 4P, N). The proximal tubules from *Ctns*^-/-^ rats at 9-months of age showed signs of degeneration and sloughing, with poorly formed brush borders (Figure 4Q). Severely affected *Ctns*^-/-^ glomeruli at 9-months of age were collapsed and contained barely recognisable podocytes that were effaced, in comparison to WT control glomeruli (Figure 4R and 4S).

### *Ctns*^-/-^ rat kidneys show loss of Lrp2 and high levels of Havcr1 and p62

Next, we immunostained kidney sections from *Ctns*^-/-^ rats at 3, 6, 9 and 12-months of age for Lrp2 (Megalin), as this proximal tubule marker is known to be down-regulated in cystinotic kidneys.^10^ At 3-months of age, the distribution of Lrp2 in *Ctns*^-/-^ kidneys was similar to that seen in wild-type animals (Figure 5B and data not shown). However, a progressive loss of Lrp2 staining was seen in *Ctns*^-/-^ kidneys from 6-months of age onwards, with only tubules near the juxtamedullary junction retaining Lrp2 at 12- and 17-months of age (Figure 5A-E and data not shown).

**Figure 5.**
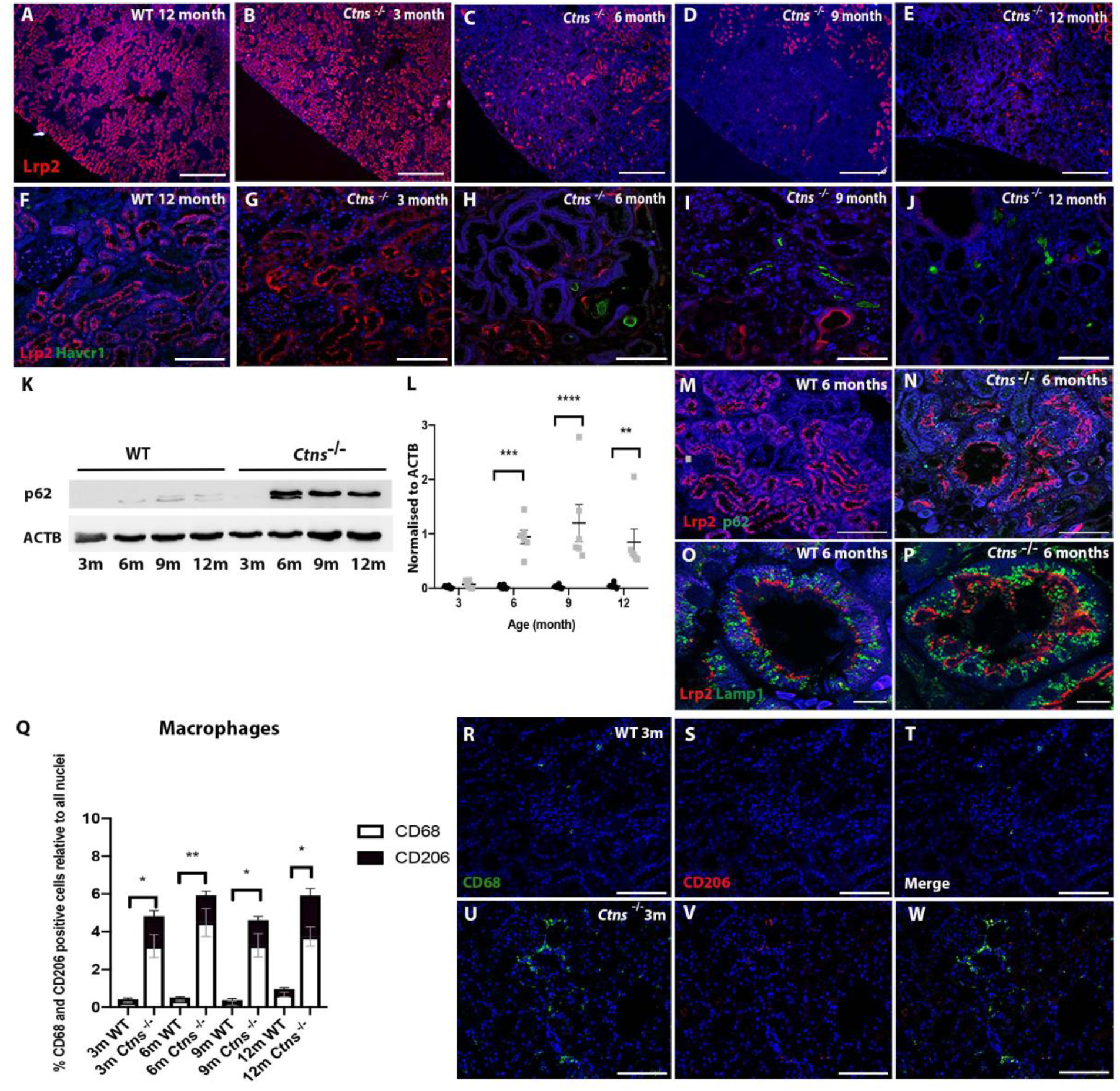
*Ctns*^-/-^ rats display progressive loss of proximal tubule marker Lrp2, increased expression of Havcr1 and autophagy pathway defect. **A-E)** Representative images of fluorescent staining with proximal tubule marker, Lrp2 in WT and *Ctns*^-/-^ rat at different timepoints. Scale bar 100 μm. **F-J)** Representative images of fluorescent staining with kidney injury marker, Havcr1 and Lrp2 in WT and *Ctns*^-/-^ rat at different timepoints. Scale bar 100 μm. **K)** Representative western blot against p62 in kidney tissue at different timepoints in WT and *Ctns*^-/-^ rats, and **L)** quantification of three independent experiments. Two-way ANOVA performed. **P<0.01, ***P<0.001, ****P<0.0001. n = 4 *Ctns*^-/-^ and 4 WT **M, N)** Representative images of fluorescent staining with autophagosome marker, p62 and Lrp2 in WT and *Ctns*^-/-^ rat at 6 months. Scale bar 100 μm. **O, P)** Representative images of fluorescent staining with lysosomal marker, Lamp1 and Lrp2 in WT and *Ctns*^-/-^ rat at 6 months. Scale bar 10 μm. Nuclei counterstain in (A-J) and (M–P) was Hoechst. **Q)** Graph displaying percentage of CD68 and CD206 positive cells relative to the number of nuclei in WT and *Ctns*^-/-^ rat at different timepoints. Two-way ANOVA performed. *P<0.05, **P<0.01. **R-W)** Representative images of fluorescent staining with pan macrophage marker CD68 and M2 marker CD206 in WT and *Ctns*^-/-^ rat at 3-months. Scale bar 100 μm.

To determine if this loss of Lrp2 was preceded by injury, we immunostained *Ctns*^-/-^ and control kidneys at 3-months of age for Havcr1, a specific and sensitive marker of proximal tubule damage. However, we observed little to no Havcr1 staining in either *Ctns*^-/-^ or control kidneys at 3-months of age (Figure 5G and data not shown). Instead, Havcr1^+^ cells became more readily apparent in *Ctns*^-/-^ kidneys at 6-months of age in tubular casts (Figure 5H). At 9-months of age, Havcr1 staining is seen more widely on the apical surface of *Ctns*^-/-^ tubules that had down-regulated Lrp2 (Figure 5I). At 12-months of age, we found Havcr1 staining in *Ctns*^-/-^ kidneys to be largely restricted to occasional interstitial cells and remnant tubules (Figure 5J). No Havcr^+^ cells were detected in control kidneys at any timepoint (Figure 5F and data not shown).

We, and others, have shown that cystinotic cells have autophagy flux defects and high numbers of autophagosomes.^26,27^ To examine this in *Ctns*^-/-^ rats, we performed western blot analysis for the autophagy marker SQSTM1/p62 (herein referred to as p62) on whole kidney lysates at 3-, 6-, 9- and 12-months of age. We observed a significant increase in p62 levels at 6-, 9- and 12-months of age in *Ctns*^-/-^ rat kidneys compared to WT littermate controls, consistent with a greater number of autophagosomes in *Ctns*^-/-^ tubules (Figure 5K and 5L). This result was confirmed by immunostaining kidney sections for p62 and Lrp2, showing a qualitative increase in the number of p62^+^ puncta in the proximal tubules of 6-month old *Ctns*^-/-^ rats (Figure 5M and 5N). We also stained these sections for the lysosomal marker Lamp1, revealing a greater number and size of lysosomes in *Ctns*^-/-^ Lrp2^+^ proximal tubules at 6-months of age compared to controls (Figure 5O and P).

To further explore the inflammatory response revealed by the urine and histological analyses, we co-immunostained *Ctns*^-/-^ and control kidneys at 3, 6, 9 and 12-months of age with CD68, a pan macrophage marker and CD206, a marker of anti-inflammatory macrophages (M2). Quantification of CD68^+^ cells revealed significantly more macrophages/monocytes in *Ctns*^-/-^ kidneys compared to WT kidneys at all timepoints with approximately a third to a half of the cells being positive for both CD68 and CD206 (Figure 5Q; representative staining’s of *Ctns*^-/-^ and control kidneys at 3 months of age are shown in Figure 5 R-W).

### Behavioural and bone phenotypes of *Ctns*^-/-^ rats

The *Ctns*^-/-^ mouse is reported to be less active compared to WT littermates.^8^ To determine if *Ctns*^-/-^ rats display a similar behavioural defect, *Ctns*^-/-^ and control rats were assessed at 9- and 12-months of age. An open field apparatus was used to analyse locomotive activity and exploratory behaviour in a familiar environment. No significant differences were observed between *Ctns*^-/-^ and control rats of either sexfor total activity, total time spent in the inner part of the arena or total rearing behaviours at either timepoint (Supplementary Figure 1G-L).

As bone defects are also characteristic of cystinosis,^8^ we undertook an X-ray examination of *Ctns*^-/-^ and control rats at 12- and 17-months of age. No gross skeletal abnormalities were detected in either males or females (data not shown). However, microCT analysis revealed that female *Ctns*^-/-^ animals at 17-months of age show a significant reduction in cortical bone cross-sectional thickness, bone area, and trabecular thickness compared to WT littermates (Supplementary Table 1 and Supplementary Figure 1M-T). There were insufficient numbers of males at 17-months of age to collect meaningful data; however there was a significant reduction in cortical tissue mineral density, trabecular thickness and an increase in connectivity density in *Ctns*^-/-^ males at 12-months of age (Supplementary Table 1).

## Discussion

In this report we describe the characterisation of a new rat model of cystinosis generated on the SD background using gene editing. These rats accumulate cystine, including crystal deposits, in multiple tissues from 3 months of age and develop Fanconi syndrome at 3-6 months of age, as defined by polyurea, polydipsia, and excessive urinary loss of protein, glucose, calcium and albumin. The Fanconi syndrome is more severe at 9-months of age, with pronounced urinary loses of phosphate, urea and creatinine, together with significantly elevated plasma creatinine indicative of reduced GFR and renal failure. We found that plasma urea levels, which are a less reliable indicator of kidney function, are not elevated in *Ctns*^-/-^ rats until 17-months of age. This late rise in plasma urea likely reflects the severe degree of renal failure at this stage as in humans, GFR must be reduced by ~50% before plasma urea levels become abnormal.^28^

The onset of kidney dysfunction in *Ctns*^-/-^ rats is earlier than that reported for C57BL/6 *Ctns*^-/-^ mice, which develop a mild/incomplete Fanconi syndrome phenotype between 2-9 months of age with no serum abnormalities until 10-18-months of age, coinciding with the start of renal failure.^9–12^ In humans, symptoms of renal Fanconi syndrome usually become apparent between 6-12 months of age and a gradual decrease in GFR starts during childhood. Thus, the timing of the renal defects in our cystinotic rat model closely recapitulates the progression of the disease seen in humans.^13,29^

Histologically, *Ctns*^-/-^ rats begin to develop pathological lesions from 3-months of age that include both glomerular and tubular abnormalities. The presence of glomerular defects in *Ctns*^-/-^ rats is notable as these are not particularly pronounced in other animal models of cystinosis. Indeed, multinucleated podocytes, which are observed in our cystinotic rats from 9-months of age onwards, have only been reported previously in humans.^30^ In human patients, renal histopathology begins with flattening of proximal tubule cells and apoptosis between 6-12 months of age, leading to the characteristic ‘swan neck’ lesion.^31,32^ Much like the human scenario we find the ‘swan neck’ lesion first appears in *Ctns*^-/-^ rats at 3-months of age and coincides with the manifestation of Fanconi syndrome. By contrast, in the C57BL/6 mouse model of cystinosis the ‘swan neck’ lesion is not readily observed until 6-months of age.^12,33^

As noted previously in humans and other animal models of cystinosis,^10,34,35^ we found a progressive loss of Lrp2 in the superficial kidney cortex regions of *Ctns*^-/-^ rats, starting at 6-months of age. These Lrp2-deficient proximal tubule cells, corresponding to the S1 and S2 segments, will be compromised in receptor-mediated endocytosis and this will be a major contributor to the renal Fanconi syndrome.^36^ It has been suggested that the decrease in Lrp2 in the S1/S2 segments is an adaptive dedifferentiation response that serves to protect these cells from proteolytic pressure caused by a compromised endo-lysosomal system.^12^ We found that the proximal tubule injury/cell death marker Havcr1 was upregulated in cells lacking Lrp2, consistent with these cells being damaged and/or in the process of dying. Contributing to proximal tubule cellular stress may be an impairment in basal autophagy and an accumulation of autophagosomes, which we characterised in prior work using cystinotic induced pluripotent stem cells and kidney organoids.^27^ In support of this, we observed an accumulation of the autophagosome marker p62 in *Ctns*^-/-^ rat kidney tissue, similar to that reported in human kidneys.^26^ Further work is needed to determine how much of a contribution the block in autophagy makes to the cellular stress of cystinotic S1/S2 proximal tubule cells. It is interesting to note however, that conditional knockout of *Atg7*, a key autophagosome forming enzyme, in mouse renal proximal tubular cells leads to an increase in p62, Havcr1 and apoptosis in an age-dependent manner (Suzuki et al., 2019).^36^ As autophagy can be protective against acute kidney injury (AKI) under certain contexts, therapeutic approaches that activate autophagy may be beneficial for individuals with cystinosis.^38^

We found that *Ctns*^-/-^ rat kidneys still retain Lrp2 staining at 17-months of age in the inner cortex corresponding to the location of the S3 segment of the proximal tubule. This suggests that the S3 segment is more resilient to cystine accumulation and may be conferring sufficient renal resorptive/secretory function to support survival in the face of compromised S1/S2 function. This resilience of the proximal tubules of the outer medullary region may be related to the more unhospitable environment of this part of the kidney, namely low blood flow, with these cells being more adapted to handle cellular stress. Indeed, regarding autophagy, deletion of *Atg5* specifically in S3 cells is not detrimental to renal outcomes following AKI and instead shows some benefits, suggesting that injured S3 cells are not dependent on autophagy for their survival.^39^ It will be interesting to explore in the future whether there is a progressive loss of S3 cells or function in *Ctns*^-/-^ rats with aging that correlates with the gradual decline in GFR, as this will help focus therapeutic efforts on preserving S3 survival.

We noted a recruitment of macrophages into the kidney of *Ctns*^-/-^ rats at 2-months of age prior to the onset of Fanconi syndrome, making the immune response one of the earliest features of the disease. Prior work has shown that blocking macrophage recruitment in cystinotic mice leads to improvements in kidney function and structure and is consistent with the notion that stressed proximal tubule cells recruit immune cells and establish a chronic inflammatory state that helps drive renal damage.^40^ With our *Ctns*^-/-^ rat model now in hand, it will be possible in follow-on studies to examine the effect of early immunosuppression on the progression of the disease.

In summary, we present here a new rat model of cystinosis that faithfully recapitulates the human disease in terms of timing, severity and histopathological changes. In this regard, *Ctns*^-/-^ rats are superior to other existing animal models and will be a vital tool to further our understanding of cystinosis and develop new therapies that slow or stop the progression of the disease.

## Acknowledgements

We thank the following people for their help; Adrian Turner, Jacqui Ross, Praju Vikas Anekal and Ratish Kurian for electron microscopy and microscopy. Satya Amirapu for histology. Dr Jordi Boix-i-Coll for behavioural studies. Vernon Jansen Unit for animal husbandry.

## Authors contributions

Dr. Hollywood and Prof. Davidson conceptualized the study, wrote the manuscript and acquired funding; Drs Hollywood, Davidson and Lewis designed experiments and performed analysis; Dr Hollywood, Mr. Sreebhavan, Ms Cheung, Ms Chatterjee, Drs Martis, Buckel and Mathews performed experiments; Dr Kallingappa established cystinotic rat lines. All authors reviewed manuscript.

---

**Supplementory Table 1:** Microarchitectural parameters of tibias from WT and Ctns^-/-^ rats as determined by microCT. Unpaired t-test performed. Values are means ± SEM. * p<0.05 vs. WT.

|  | 12 month |  | 17 month |  |
| --- | --- | --- | --- | --- |
| **Females** | WT (n=5) | *Ctns*<sup>-/-</sup> (n=9) | WT (n=5) | *Ctns*<sup>-/-</sup> (n=6) |
| **Tibia length (mm)** | 41.5 ± 0.5 | 40.7 ± 0.6 | 42.6 ± 0.3 | 40.5 ± 1.4 |
| **Proximal tibia trabecular bone** |  |  |  |  |
| **Bone volume fraction (%)** | 20.2 ± 2.3 | 19.6 ± 1.8 | 23.4 ± 2.0 | 17.7 ± 3.4 |
| **Trabecular thickness (μm)** | 78.6 ± 3.5 | 72.2 ± 2.2 | 102.5 ± 3.7 | 83 ± 2.8* |
| **Trabecular separation ($\mu\text{m}$)** | 257 ± 18 | 267 ± 13 | 311 ± 37 | 448 ± 95 |
| **Trabecular number (1/mm)** | 2.55 ± 0.21 | 2.69 ± 0.19 | 2.32 ± 0.29 | 2.11 ± 0.37 |
| **Connectivity density (1/mm<sup>3</sup>)** | 100.6 ± 12.1 | 139.4 ± 12.0* | 50.8 ± 16.1 | 53.5 ± 10.5 |
| **Tibia cortical bone** |  |  |  |  |
| **Cross-sectional tissue area (mm<sup>2</sup>)** | 7.1 ± 0.2 | 6.3 ± 0.3 | 7.3 ± 0.3 | 6.3 ± 0.3* |
| **Cross-sectional bone area (mm<sup>2</sup>)** | 4.9 ± 0.1 | 4.3 ± 0.2 | 5.3 ± 0.2 | 3.9 ± 0.2* |
| **Cortical thickness ($\mu\text{m}$)** | 439.8 ± 19.7 | 409.9 ± 14.2 | 524.0 ± 12.0 | 399.1 ± 20.1* |
| **Medullary area (mm<sup>2</sup>)** | 2.2 ± 0.2 | 2.0 ± 0.2 | 2.0 ± 0.1 | 2.4 ± 0.2 |
| **Cortical tissue mineral density (g/cm<sup>3</sup>)** | 1.348 ± 0.009 | 1.343 ± 0.008 | 1.320 ± 0.324 | 1.341 ± 0.013 |
| **Polar moment of inertia (mm<sup>4</sup>)** | 10.1 ± 0.2 | 7.7 ± 0.7* | 11.0 ± 0.8 | 7.4 ± 0.6* |
|  | **12 month** |  |  |  |
| **Males** | WT (n=6) | *Ctns*<sup>-/-</sup> (n=8) |  |  |
| **Tibia length (mm)** | 48.7 ± 0.6 | 48.6 ± 0.9 | | |
| **Proximal tibia trabecular bone** |  |  |  |  |
| **Bone volume fraction (%)** | 11.1 ± 0.6 | 10.7 ± 1.4 | | |
| **Trabecular thickness ($\mu\text{m}$)** | 96.2 ± 5.1 | 78.9 ± 3.3* | | |
| **Trabecular separation ($\mu\text{m}$)** | 530.0 ± 35.1 | 479.4 ± 59.4 | | |
| **Trabecular number (1/mm)** | 1.17 ± 0.10 | 1.36 ± 0.17 | | |
| **Connectivity density (1/mm<sup>3</sup>)** | 24.5 ± 4.4 | 47.1 ± 8.2* | | |
| **Tibia cortical bone** |  |  |  |  |
| **Cross-sectional tissue area (mm<sup>2</sup>)** | 13.3 ± 0.5 | 12.7 ± 0.7 | | |
| **Cross-sectional bone area (mm<sup>2</sup>)** | 7.6 ± 0.1 | 7.1 ± 0.3 | | |
| **Cortical thickness ($\mu\text{m}$)** | 483.1 ± 17.8 | 452.4 ± 16.0 | | |
| **Medullary area (mm<sup>2</sup>)** | 5.7 ± 0.5 | 5.6 ± 0.5 | | |
| **Cortical tissue mineral density (g/cm<sup>3</sup>)** | 1.319 ± 0.003 | 1.310 ± 0.002* | | |
| **Polar moment of inertia (mm<sup>4</sup>)** | 29.2 ± 1.4 | 26.4 ± 2.2 | | |

## Legends to figures

**Supplementary Figure 1.**
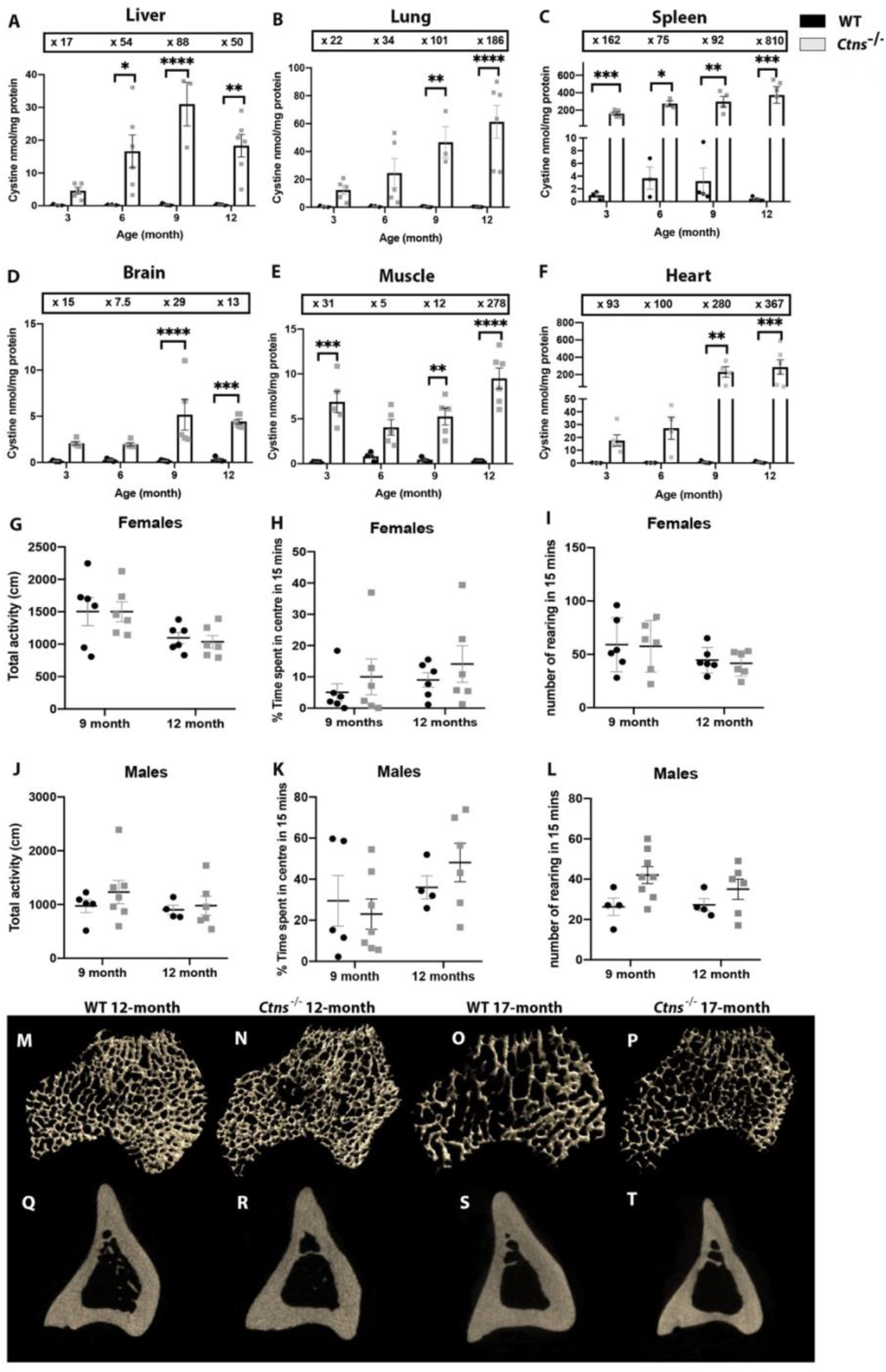
**A-F)** Amount of cystine (nmol/mg of protein) in WT and *Ctns*^-/-^ rats at different timepoints in various tissues. Boxed values indicate fold-change compared to WT at same timepoint. Two-way ANOVA performed, all data are plotted mean ± SEM. *P<0.05, **P<0.01, ***P<0.001. n = 5 WT and 6 *Ctns*^-/-^ per group. **G-L)** Graphs displaying quantification of total activity (cm), percentage of time spent in the inner area of the arena and number of rearing behaviours in 15 mins in both females and males. Two-way ANOVA performed, no significance observed. n= 13 *Ctns*^-/-^ and 11 WT. Representative images of trabecular **(M-P)** and cortical **(Q-T)** bone in tibias from female WT and *Ctns*^-/-^ rats at 12 and 17 months of age. n= 14 *Ctns*^-/-^ and 13 WT.

